# DNA methylation of PGC-1α is associated with elevated mtDNA copy number and altered urinary metabolites in Autism Spectrum Disorder

**DOI:** 10.1101/2021.01.20.427429

**Authors:** Sophia Bam, Erin Buchanan, Caitlyn Mahony, Colleen O’Ryan

## Abstract

**Background:** Autism Spectrum Disorder (ASD) is a complex disorder that is underpinned by numerous dysregulated biological pathways, including canonical mitochondrial pathways. Epigenetic mechanisms contribute to this dysregulation and DNA methylation is an important factor in the aetiology of ASD. We examined the relationship between DNA methylation of peroxisome proliferator-activated receptor gamma coactivator-1 alpha (PGC-1α), an essential transcriptional regulator of mitochondrial homeostasis, and mitochondrial dysfunction in an ASD cohort of South African children.

**Results:** Using targeted Next Generation bisulfite sequencing, we found 12 highly variable CpG sites in PGC-1α that were significantly differentially methylated (p<0.05) between ASD (n = 55) and controls (n = 44). In ASD, eight CpG sites were hypermethylated in the PGC-1α promotor with a putative binding site for CAMP response binding element 1 (CREB1) spanning one of these CpG sites (p = 1 × 10^−6^). Mitochondrial DNA (mtDNA) copy number, a marker of mitochondrial function, was elevated (p = 0.002) in ASD compared to controls and correlated significantly with DNA methylation at the PGC-1α promoter. There was a positive correlation between methylation at PGC-1α at CpG#1 and mtDNA copy number (Spearman’s r = 0.2, n = 49, p = 0.04) in ASD, but a negative correlation between methylation at PGC-1α at CpG#4 promoter and mtDNA copy number in controls (Spearman’s r = −0.4, n = 42, p = 0.045). While there was no relationship between mtDNA deletions and PGC-1α methylation in ASD, mtDNA deletions correlated negatively with methylation at PGC-1α at CpG#4 (Spearman’s r = −0.4, n = 42, p = 0.032) in controls. Furthermore, levels of urinary organic acids associated with mitochondrial dysfunction correlated significantly (p<0.05) with DNA methylation at PGC-1α CpG#1 and mtDNA copy number in ASD (n= 20) and controls (n= 13) with many of these metabolites involved in altered redox homeostasis and neuroendocrinology.

**Conclusions:** These data show an association between PGC-1α promoter methylation, elevated mtDNA copy number and metabolomic evidence of mitochondrial dysfunction in ASD. This highlights an unexplored link between DNA methylation and mitochondrial dysfunction in ASD.

## BACKGROUND

Autism Spectrum Disorder (ASD) is defined by the presence of behavioural traits (1) despite being a highly heritable neurodevelopmental disorder (2). ASD is characterised by deficits in social communication and restrictive, repetitive behaviours (3). ASD is a complex disorder that affects the central nervous system as well as other organ systems, implying the dysregulation of pleiotropic biological and developmental pathways. ASD is underpinned by a heterogeneous genetic architecture that includes rare *de novo* genetic variations and low risk, common single nucleotide mutations (4,5). There is increasing evidence for the role of epigenetic alterations, and DNA methylation in particular, in modulating ASD phenotypes. This is evident from discordant identical ASD twin studies (6), studies using brain tissue from individuals with ASD (7,8), with recent reviews collating numerous reports on ASD epigenetics (9–12).

Given the varied phenotypes and co-morbidities observed in individuals with ASD, numerous and diverse biological pathways have been implicated in ASD aetiology. These include gene regulatory-, signaling-, synaptic-, and mitochondrial**-**pathways (13-16). Mitochondrial dysfunction is emerging as a key contributor to ASD aetiology; deficiencies in oxidative phosphorylation (OXPHOS) can decrease the production of adenosine 5’-triphosphate (ATP), which is an essential requirement for brain function and neurodevelopment. The observation that congenital errors of mitochondrial metabolism contribute to > 5% of ASD cases (17) first implied a role for mitochondrial dysfunction in ASD. This has since been supported by clinical (18), biochemical (19,20), molecular (21) and more recently, epigenetic data (22). Citrigno *et al* (23) comprehensively reviewed the recent experimental data that support the Mitochondrial Dysfunction Hypothesis in ASD.

Mitochondrial homeostasis is dynamic and is regulated by interdependent pathways that govern mitogenesis, mitophagy, mitochondrial fission and fusion (24). These dynamic mechanisms enable cellular adaptation to changing energy demands, nutrient availability and oxidative stress by regulating mitochondrial DNA (mtDNA) copy number (25,26). Abnormal or fluctuating levels of mtDNA copy number is a marker of mitochondrial dysfunction (27,28). An essential transcriptional regulator of mitochondrial homeostasis is peroxisome proliferator-activated receptor gamma coactivator-1 alpha (PGC-1α) which regulates fatty acid ,*β*-oxidation, OXPHOS, gluconeogenesis and antioxidant defense responses (29). PGC-1α catalyses mitogenesis by upregulating nuclear respiratory factors 1 and 2 (Nrf-1 and Nrf-2), which promotes the transcription of mitochondrial transcription factors A (TFAM) and B2 (TFB2M) (24). Importantly, numerous studies report a correlation between DNA methylation of the PGC-1α promoter and PGC-1α transcription, mtDNA copy number and metabolic disease (30–33), suggesting that DNA methylation regulates PGC-1α-driven mitogenesis. Changes in mitochondrial morphology via fission and fusion, together with mitogenesis, also contribute to maintaining mitochondrial metabolism and function by limiting reactive oxygen species (ROS) damage (34). Mitochondrial fusion occurs in response to mild oxidative stress and is mediated by mitofusins 1 and 2 (MFN1 and MFN2) and optic atrophy 1 (OPA1) (35– 37). These proteins work in conjunction with accessory proteins, such as stomatin-like protein 2 (STOML2), which maintains the long isoforms of OPA1 needed to fuse the inner mitochondrial membranes (38). Fission occurs during severe oxidative stress and separates damaged mitochondrial components from the healthy mitochondrial network. Fission is mediated by dynamin-related protein 1 (DRP1) which works with mitochondrial fission protein 1 (FIS1) to divide the outer mitochondrial membrane (39,40). Recently, genes involved in mitogenesis, fission and fusion have been implicated in neuropathology. PGC-1α is reported to play an important role in excitatory neurotransmitter signaling, neuroprotection, neuroinflammation and neurogenesis and has been implicated in bipolar disorder, Parkinson’s disease, Huntington’s disease, schizophrenia and Alzheimer’s disease (41). In addition, recent studies have implicated PGC-1α (29), fusion and fission genes (42,43) in ASD aetiology (29).

In view of the evidence implicating both differential DNA methylation and mitochondrial dysfunction in ASD, we examined the relationship between these two processes in a cohort of South African children with ASD. First, we first determined whether PGC-1α was differentially methylated (DM) between ASD and matched controls from a South African population. Secondly, we addressed whether DNA methylation changed mitochondrial function, measured using mtDNA copy number and urinary metabolomics. Our aim was to test the hypothesis that DNA methylation of mitochondrial biogenesis genes changes mitochondrial function, thereby contributing to the aetiology of ASD.

## RESULTS

### ASD cohort phenotype and demography

The individuals in our study with ASD spanned the full range of developmental phenotypes observed in ASD. This is reflected in both the number of different Autism Diagnostic Observation Schedule, Second Edition (ADOS-2) Modules used for assessments and the autism severity scores (Table S1, Additional File 1). Each ADOS-2 Module is tailored to match different developmental levels, ranging from pre-verbal individuals with ASD to those with fluent speech. Although our cohort comprised of different demographic groups, demography did not correlate with any ASD trait or any molecular marker (data not shown).

### PGC-1α promoter is hypermethylated in ASD

We investigated whether DNA methylation contributed to the regulation of mitochondrial biogenesis by measuring the methylation of PGC-1α, a central transcriptional regulator of mitogenesis. We defined highly variable CpGs as those sites where the methylation range exceeded 5% across all samples in order to identify functionally significant DM genes. This threshold is consistent with *in vitro* (44) and *in vivo* (45) methylation studies. There were 12 highly variable CpG sites in PGC-1α that were significantly DM (p<0.05) between ASD and controls (n= 55 ASD, n= 43 controls) (Table S2, Additional File 1). Of these, eight CpG sites were hypermethylated in ASD and clustered around the transcriptional start site (TSS), between the 5’ untranslated region (UTR) and intron 1 (Fig. 1 a, b), while four sites, located at intron 2, intron 12, and the 3’UTR were hypomethylated in ASD (Fig. S1, Additional File 2).

**Fig 1:**
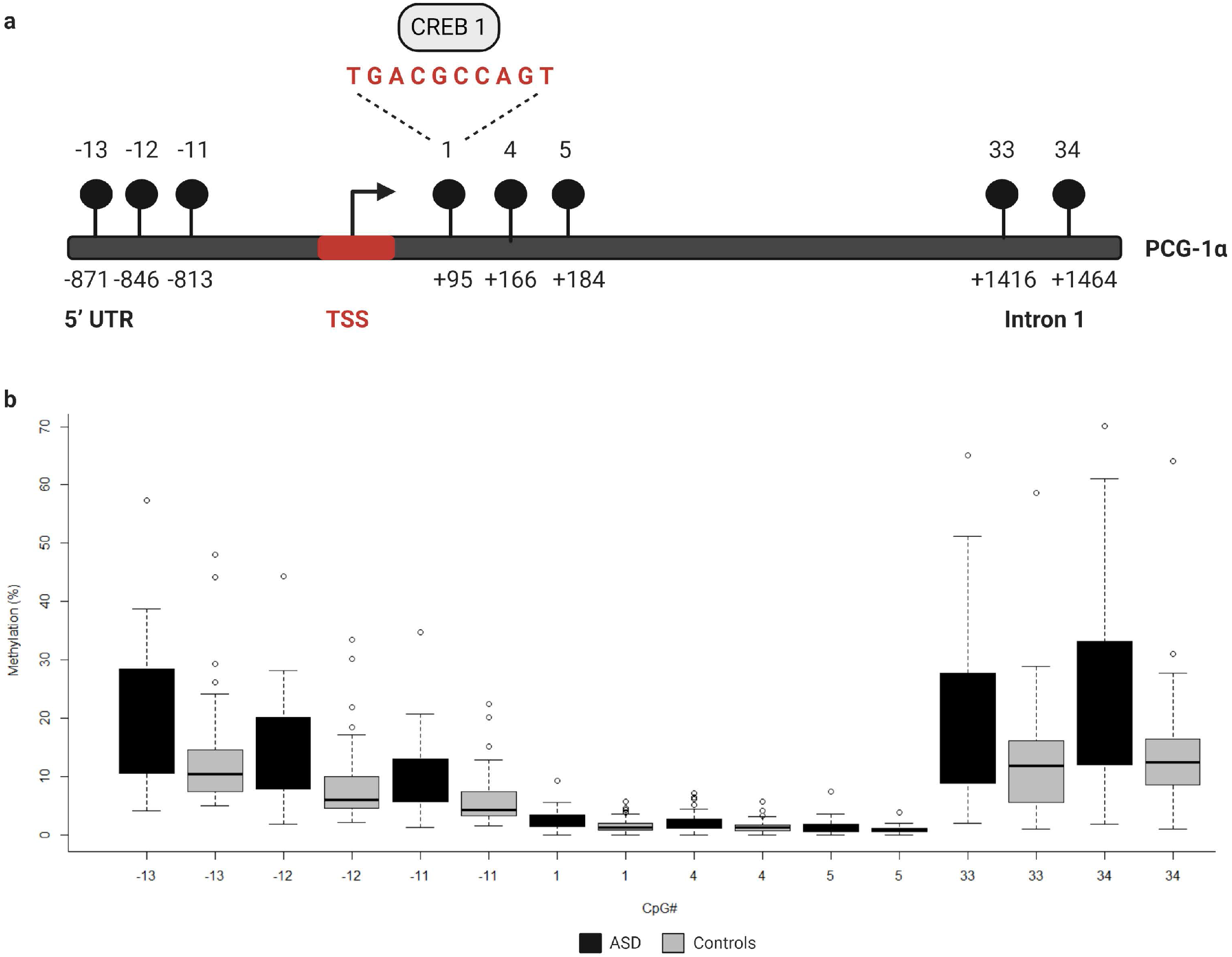
The PGC-1α promoter is hypermethylated in ASD. a) Diagrammatic representation of the PCG-1α gene promoter region (Chr4:23889974) showing the location of the eight hypermethylated CpG sites (black circles) relative to the transcription start site (TSS) and the sequence of the binding site for the transcription factor, CREB1. b) Box plot showing the percentage methylation of the eight differentially methylated PGC-1α promoter CpG sites measured using Targeted Next-Generation Bisulfite Sequencing (n = 55 ASD, n = 44 controls) (UTR = untranslated region).

The significant DM sites of PGC-1α in our study is consistent with a role for DNA methylation in regulating mitochondrial biogenesis. To examine this hypothesis, we quantified DNA methylation of nuclear respiratory factor 2 alpha subunit (NRF2A), the transcriptional regulator of mitogenesis that acts directly downstream of PGC-1α, as well as four genes involved in mitochondrial fission and fusion (STOML2, MFN2, OPA1, and FIS1) in a subset of our cohort (n= 22 ASD, n= 22 controls). STOML2 contained two DM CpG sites, located in intron 2 and exon 5 downstream of the TSS; these sites were hypermethylated in ASD (Fig. 2a; Table S2, Additional File 1). Significant DM sites between ASD and controls were also identified at one CpG site in NRF2A and FIS1, and at two sites in MFN2 and OPA1 (Fig. 2a; Table S2, Additional File 1). Therefore, we observed multiple DM mitogenesis, fission and fusion genes converging on the regulation of mitochondrial homeostasis in ASD (Fig. 2b), congruent with a role for DNA methylation in the dysregulation of mitochondrial function in our cohort.

**Fig 2:**
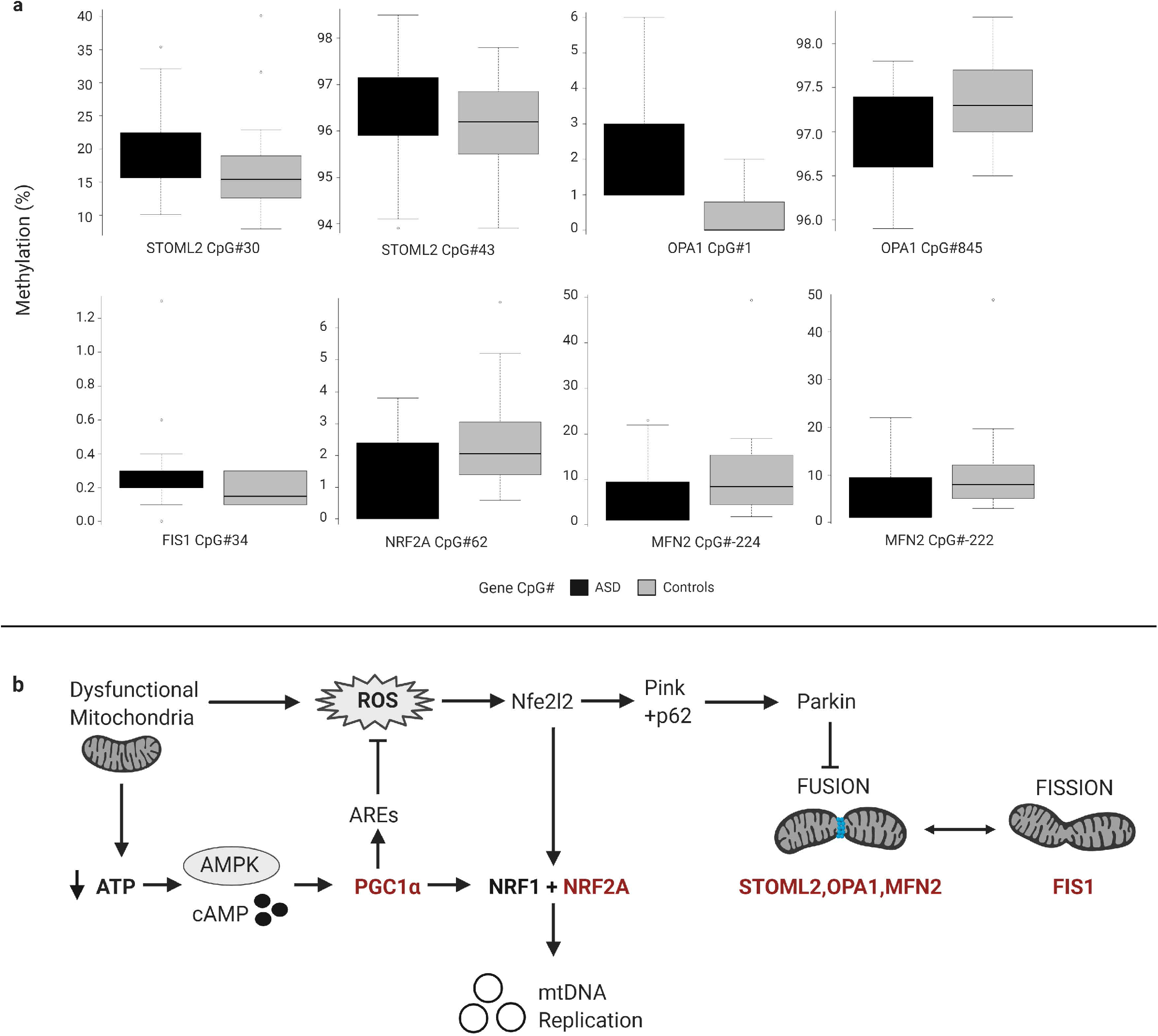
Differentially methylated genes in ASD converge on pathways regulating mitochondrial homeostasis. a) Targeted Next-Generation Bisulfite Sequencing of STOML2 (n = 55 ASD, 44 controls) and FIS, MFN2, OPA1 and NRF2A (n = 22 ASD, n = 22 controls) with differential methylation at two CpG sites in STOML2, MFN2 and OPA1 and one site in FIS1 and NRF2A. Data represent mean percentage methylation in ASD relative to controls and error bars represent sample standard deviation as a fraction of average methylation at each site. Significant differential methylation was identified using a two-tailed unpaired t-test with unequal variance (p<0.05). b) Mitochondrial homeostasis is maintained by differentially methylated genes (in red) involved in mitochondrial biogenesis, fission and fusion that converge on the regulation of mitochondrial copy number in response to metabolic and oxidative stress. Metabolic stress (decrease in ATP production) activates cAMP and AMPK signalling, leading to the transcription and activation of PGC-1α. PGC-1α upregulates the expression of Nrf1 and NRF2A which induce the transcription of the mitochondrial transcription factors to facilitate mitogenesis. PGC-1α also activates AREs to modulate oxidative stress or ROS levels. Oxidative stress activates the redox sensitive Nfe2l2 pathway, which upregulates both NRF2A and the Pink-Parkin pathway that controls mitophagy, fission and fusion. (AREs = antioxidant response elements; mtDNA = mitochondrial DNA; ROS = reactive oxygen species).

### Is PGC-1α hypermethylation associated with altered mitochondrial function?

We examined whether methylation affected mitochondrial biogenesis and function because DNA methylation was altered at several key regulators of mitogenesis in our ASD cohort. *In silico* transcription factor binding site analysis of DM CpG sites in the PGC-1α promoter predicted a putative binding site for the transcription factor CAMP response binding element 1 (CREB1) at the CpG#1site (p = 1 × 10^−6^). This suggests that the DM site in ASD may have functional significance, which we examined by assessing mitochondrial function in our cohort. This was done by quantifying mtDNA copy number and deletions, as well as the levels of urinary metabolites typically associated with mitochondrial disease in ASD relative to controls.

### Mitochondrial DNA copy number is increased in ASD

Increased mtDNA copy number is a compensatory response to mild oxidative stress (25,26) and is an established biomarker of mitochondrial function (46). Mitochondrial DNA copy number of NADH dehydrogenase 1 (ND1) relative to Beta-2-microglobulin (B2M) was significantly elevated in the ASD group compared to the control group (p = 0.002) (Fig. 3a). Mitochondrial deletions are typical of mitochondrial disease, therefore we examined whether mitochondrial deletions differed in ASD compared to controls by quantifying the copy number of the mitochondrial gene ND1 relative to NADH dehydrogenase 4 (ND4); the latter resides in the major mitochondrial deletion arc. While mtDNA deletions were not significantly elevated in ASD relative to controls (p = 0.162), we observed markedly elevated mtDNA deletions in some ASD individuals (Fig. 3b). Notably, mtDNA copy number correlated significantly (Spearman’s r = 0.9, n = 49, p = 8.814 ×10^−10^) with mtDNA deletions (Fig. 3c), which suggests that elevated mtDNA copy number is associated with mitochondrial dysfunction in our cohort.

**Fig 3:**
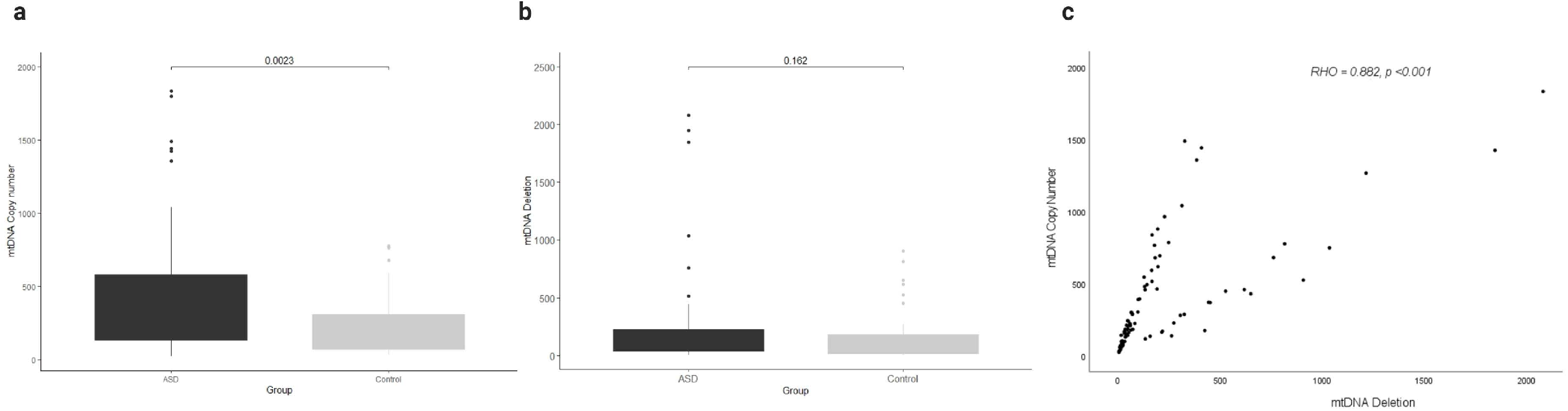
Relationship between mitochondrial DNA (mtDNA) copy number and mtDNA deletion in ASD. a) mtDNA copy number is significantly elevated in ASD. Relative quantification of mtDNA copy number was performed by multiplex real-time qPCR of ND1 and B2M (n = 59 ASD, n = 40 control). b) A non-significant increase in mtDNA deletions is observed in ASD. Relative quantification of mtDNA deletions was performed by multiplex real-time qPCR of ND1 and ND4 (n = 59 ASD, n = 40 control). Significance was established using Student t-tests where (p < 0.05). c) mtDNA copy number correlates significantly with mtDNA deletions in ASD and controls (n=99), where Spearman’s rho = 0.882, p<0.001.

### PGC-1α promoter methylation correlates significantly with mtDNA copy number and this relationship is altered in ASD

We examined whether PGC-1α promoter methylation is associated with mtDNA copy number and/or deletions, and thus mitochondrial function in ASD. We observed a significant correlation between DNA methylation at the PGC-1α promoter and mtDNA copy number, and this relationship differed between ASD and control groups. In the control group, PGC-1α methylation at CpG#4 correlated negatively with both mtDNA copy number (Spearman’s r = −0.4, n = 42, p = 0.045) and mtDNA deletions (Spearman’s r = −0.4, n = 42, p = 0.032) (Fig. S2, Additional File 3). However, in the ASD group there was a significant positive correlation between PGC-1α methylation at CpG#1 and mtDNA copy number (Spearman’s r = 0.9, n = 49, p = 0.04) (Fig. S2a, Additional File 3), with no correlation between PGC-1α methylation and mtDNA deletions This suggests that PGC-1α methylation is associated with mitochondrial biogenesis and function and that this relationship is disrupted in our ASD group.

### DNA methylation and mtDNA copy number are associated with metabolomic markers of mitochondrial dysfunction

We investigated whether the differential methylation of PGC-1α and elevated mtDNA copy number observed in ASD were associated with metabolomic evidence of mitochondrial dysfunction. We examined the correlation between PGC-1α methylation, mtDNA copy number and levels of urinary organic acids which had previously been measured (n= 20 ASD, n= 13 controls) using gas chromatography −mass spectrometry (GC-MS) (22). DNA methylation at PGC-1α CpG#1, which is DM in our samples, correlated significantly with three of the 55 urinary organic acids tested (Table S3b, Additional File 1) MtDNA copy number correlated with 22 urinary metabolites associated with mitochondrial dysfunction (Table S3a, Additional File 1). Notably, both PGC-1α methylation and mtDNA copy number are associated with metabolites derived from BCAA metabolism, including 3-hydroxy-3-methylglutaric acid (3-H-3-MGA), which correlated significantly with both PGC-1α CpG#1 (p=0.009) and with mtDNA copy number (p=0.004) (Fig. 4a, b). In addition, mtDNA copy number is associated most significantly (r< −0.3 or r> 0.3; p<0.01) with metabolites derived from four metabolic pathways: fatty acid oxidation, phenylalanine-dopamine synthesis pathway, glycine-glutamine metabolism and branched-chain amino acids (BCAAs) (Table S3a, Additional File 1). These pathways converged on mitochondrial OXPHOS, one-carbon metabolism and neurotransmitter synthesis (Fig. 5). Our data are consistent with an established metabolomic model for altered mitochondrial metabolism and neuroendocrinology (47) and supports an association between DNA methylation, elevated mtDNA copy number and mitochondrial dysfunction in ASD.

**Fig 4:**
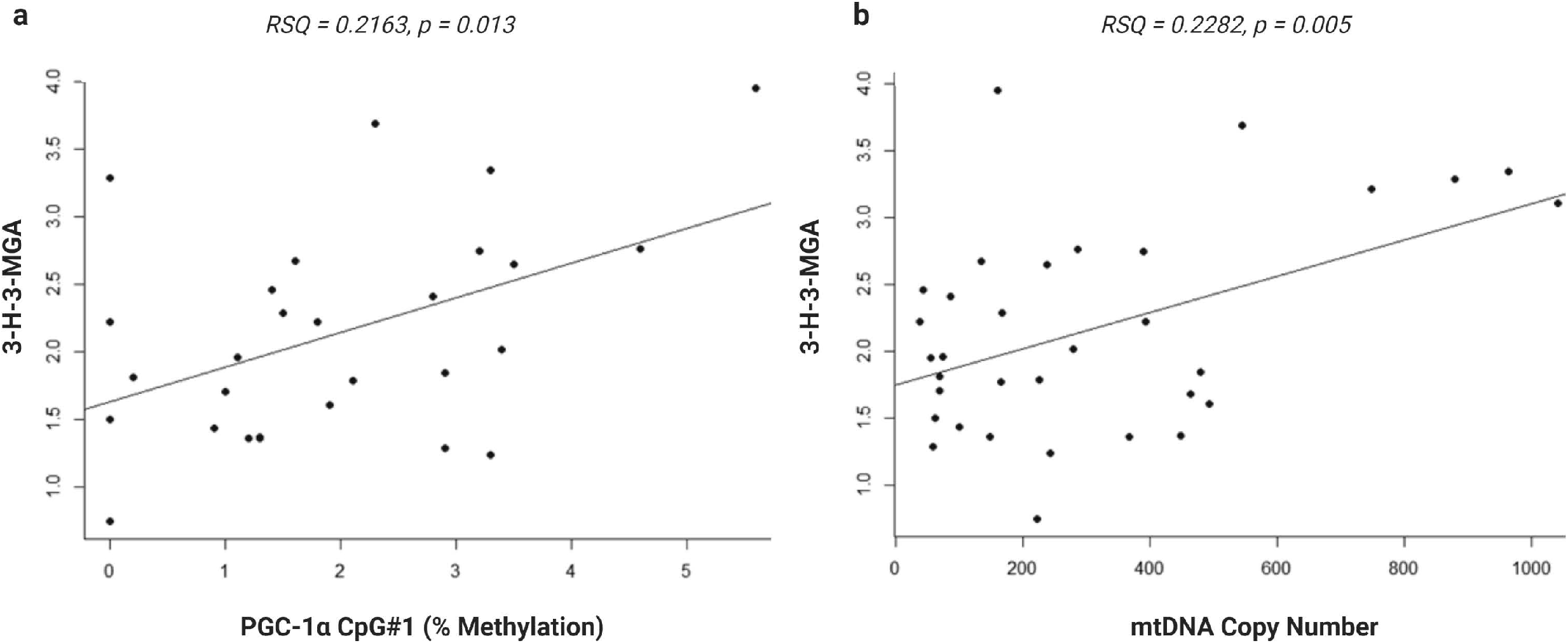
Urinary metabolite, 3-hydroxy-3-methylglutaric acid (3-H-3-MGA), correlates with PGC-1α methylation and mitochondrial DNA (mtDNA) copy number. Normal linear regression analysis shows that a) DNA methylation at PGC-1α CpG#1 correlates with 3-H-3-MGA levels, RSQ = 0,2163, Spearman Rho = 0.386, p = 0,043 and b) mtDNA copy number correlates with 3-H-3-MGA, RSQ = 0,2282, Spearman Rho= 0.355, p= 0,004; (n= 20 ASD, n= 13 controls).

**Fig 5:**
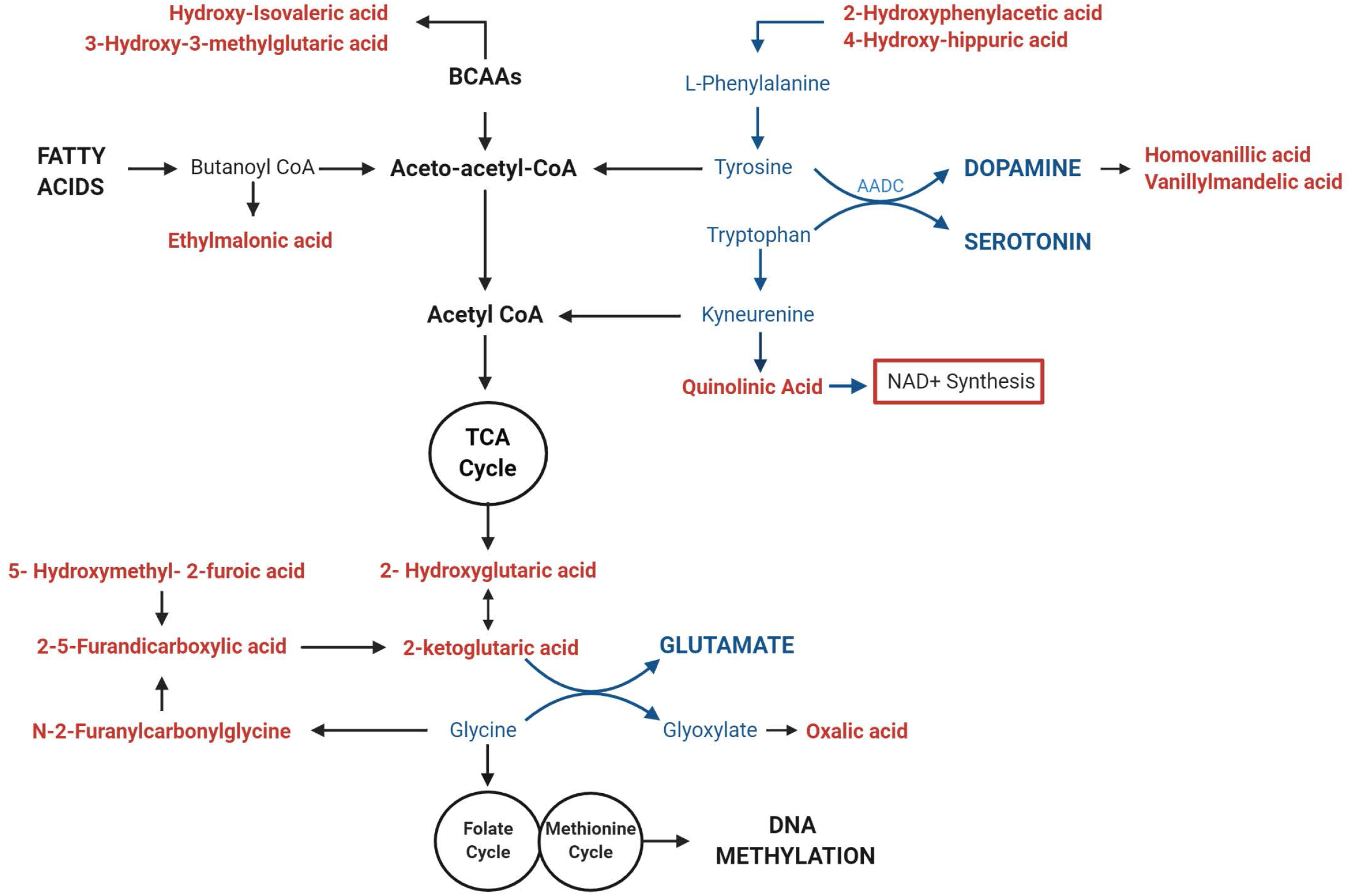
Mitochondrial DNA (mtDNA) copy number correlates with urinary metabolites implicating mitochondrial metabolism and neurotransmitter synthesis. Urinary metabolites that most significantly correlate with mitochondrial DNA copy number (p < 0.01) converge on pathways involved in the metabolism of Fatty Acids (Ethylmalonic Acid); Branched Chain Amino Acids (BCAAS) (Hydroxy-isovaleric acid, 3-hydroxy-3-methylglutaric acid); and neurotransmitters including Dopamine (2-Hydroxyphenylacetic acid, 4-Hydroxyhippuric acid, Vanillylmandelic acid and Homovanillic acid), Serotonin (Quinolinic acid), Glutamate (2-ketoglutaric acid, D-L-Hydroxyglutaric acid; 2,5-Furandicarboxylic acid, Hydroxymethyl-2-furoic acid) and Glycine (N-2Furanylcarbonylglycine, oxalic acid). Metabolites that are associated with elevated mtDNA copy number are shown in red; pathways involved in neurotransmitter metabolism are highlighted in blue.

## DISCUSSION

ASD is a heritable, complex phenotype with numerous molecular pathways contributing to its aetiology (2). Despite the high heritability of ASD, there is no single or simple genetic mutation that accounts for ASD, and the disorder is characterised by phenotypic and clinical heterogeneity. This implies that epigenetic mechanisms may be important in ASD, and DNA methylation is known to contribute to ASD aetiology (11). DNA methylation dysregulates many biological pathways in ASD, including, but not limited to, immune function (48), chromatin remodeling (49), synaptic signaling and neuronal regulation (50). Increasingly, mitochondrial biological pathways have also been implicated in ASD (18,20,21). The link between mitochondrial dysfunction and ASD is unsurprising given that efficient ATP production is essential for brain development and function. However, the relationship between DNA methylation and mitochondrial function is not fully understood. Our data examined this relationship by testing the hypothesis that mitochondrial biogenesis and fusion genes are DM between ASD and controls, and that differential methylation affects mitochondrial function in ASD.

The transcriptional regulator of biogenesis, PGC-1α, was significantly DM between ASD and controls in our cohort, with the promoter region being hypermethylated in ASD. The promoter region included a CpG site (CpG#1) containing a putative transcription binding site for CREB1, which is a potent activator of PGC-1α transcription (51). Although we were not able to examine PGC-1α transcription in our cohort, this CREB1 site is reported to be DM in metabolic disease (31) which suggests that DM sites at the PGC-1α promoter region and TSS in our ASD cohort could affect gene transcription and subsequently, mitochondrial biogenesis and function.

Mitochondria are adaptive to changing cellular metabolic demands, thus they are dynamic organelles that are remodeled by biogenesis, fission and fusion (52). Therefore, we measured the methylation of additional genes involved in mitochondrial biogenesis, fission and fusion and found that NRF2A, which facilitates mtDNA replication downstream of PGC-1α in ASD, was DM (24). We also found that genes involved in mitochondrial fission (STOML2, MFN2, OPA1) and fusion (FIS1) were DM in ASD. Collectively, these DM genes converge on the pathways regulating mitochondrial homeostasis in response to metabolic and oxidative stress (Fig 2). Of note, STOML2 was hypermethylated at two CpG sites downstream of the TSS in ASD. STOML2 plays an important role in mitochondrial fusion by stabilizing the OPA1 protein, which facilitates fusion of the inner mitochondrial membranes (36–38). STOML2-deficient cells fail to undergo mitochondrial fusion during stress, leading to mitochondrial fragmentation (38). STOML2 is also well-established as an anti-apoptotic gene in cancer cells, highlighting the importance of fusion to re-establish mitochondrial homeostasis and prevent mitophagy (autophagy of mitochondria) under stress. Our data is consistent with previous work showing that both fission and fusion genes are differentially expressed in ASD (20,42,43). This supports the link between ASD and mitochondrial fusion and fission which highlights the differential methylation of mitochondrial genes on an integrated pathway level.

Our data show genes that regulate mitochondrial biogenesis, fission and fusion are DM in our cohort. While we were unable to measure gene expression because of the low integrity of RNA extracted from the tissue source (buccal swabs) used in our study, previous studies have shown that differential methylation alters the expression of these genes (20,42,43,53). We investigated whether mitochondrial DNA copy number, a marker of mitochondrial function (46), was altered in our cohort. Changes in mtDNA copy number have been reported in ASD, with both increases (54–56) and decreases (57,58) observed in ASD. These discrepancies can be attributed to several factors that differed across studies, including the age of the participants studied, the presence of co-morbidities and the degree of mitochondrial dysfunction in participants. We observed a significant increase in mtDNA copy number in our ASD group compared to controls, which represents an established compensatory mechanism in response to mitochondrial dysfunction. We also observed a significant positive correlation of mtDNA copy number with mtDNA deletions in ASD, suggesting that elevated mtDNA copy number is indicative of mitochondrial dysfunction in our cohort. Increased mtDNA copy number is observed as a response to oxidative stress in animal models and *in vitro* studies (59–61), as well as in human clinical studies using buccal samples (62). This compensatory mechanism has also been reported in both mitochondrial diseases (63–65), neuropsychiatric and neurodevelopmental disorders (66–69).

A connection between PGC-1α methylation and mtDNA copy number is established, and a negative correlation is reported between PGC-1α promoter methylation and both PGC-1α transcription and mtDNA copy number in neurological (53) and metabolic disorders (31,33,70). Consistent with this, we report a negative correlation between PGC-1α promoter hypermethylation and both mtDNA copy number and deletions in our control group. Thus, hypomethylation of PGC-1α promoter is associated with elevated mtDNA copy number and deletions, suggesting that this is an adaptive mechanism to upregulate PGC-1α dependent mitochondrial biogenesis under conditions of mild mitochondrial dysfunction. However, this relationship between PGC-1α promoter methylation and mtDNA copy number was pertubated in ASD. We observed a significant positive correlation between PGC-1α promoter methylation at CpG#1 and mtDNA copy number. Hypermethylation at the PGC-1α promoter could inhibit the PGC-1α-dependent activation of antioxidant response elements (71) which is congruent with the evidence for elevated oxidative stress in ASD (72). Oxidative stress can induce mitogenesis via Nrf-2 (73) which upregulates NRF2A independently of PGC-1α in a redox-sensitive manner (62). We found no relationship between PGC-1α methylation and mtDNA deletions in ASD, suggesting an absence of the adaptive hypomethylation to compensate for metabolic and putative oxidative stress.

To further explore the link between differential methylation, mtDNA copy number and mitochondrial function, we used metabolomic analysis, which directly reflects the biochemical activity, including mitochondrial activity, of a biological sample (74). We examined whether differential methylation and elevated mtDNA copy number correlated with metabolomic evidence of mitochondrial dysfunction in our cohort, using urinary organic acids that had been associated with mitochondrial disease in South Africans (75). We found a metabolomic profile associated with mtDNA copy number that was consistent with a link between DNA methylation, mitochondrial dysfunction and neuropathology in our ASD cohort. Although the metabolomic analysis was performed on a smaller sample size than the methylation and mtDNA copy number experiments, it was a functional and exploratory way to corroborate the mitochondrial dysfunction reflected by elevated mtDNA copy number in our cohort. Consequently, we focused on the metabolic pathways, rather than specific metabolites, that correlated most significantly with mtDNA copy number (p<0.01) in our cohort. Metabolites derived from tyrosine, tryptophan and glycine significantly (p<0.01) correlated with mtDNA copy number. These are precursors of dopamine, serotonin, melatonin and glutamate synthesis, which are all important neurotransmitters implicated in ASD aetiology (76). The notable enrichment of metabolites derived from the dopamine pathway, which is closely tied to cellular oxidation state (77), is consistent with altered redox homeostasis in ASD. Moreover, glycine serves as the precursor to one-carbon metabolism (cysteine, methionine, and glutathione pathways). These are essential regulators of oxidative state which have been implicated in the metabolomic profile associated with ASD severity in two independent cohorts (78,79) and have been identified as a link between DNA methylation and mitochondrial dysfunction (80,81). In addition, both PGC-1α methylation and mtDNA copy number correlate significantly with metabolites derived from BCAA catabolism. BCAAs provide nitrogen to the glutamate-glutamine cycle and have been implicated as important regulators of glutamatergic neurotransmission (82), which contribute to ASD aetiology (83). Two metabolites (3-H-3-MGA and 3-methylglutaconic acid (3-MGA)) were significantly elevated in our ASD cohort (22) and are characterized as urinary biomarkers of mitochondrial respiratory chain deficiencies (75,84). Altered BCAA metabolism has also been linked to oxidative stress resulting from perturbed NAD^+^/NADH redox ratios (85). Of note, mtDNA copy number correlated significantly with the direct precursor of *de novo* NAD^+^ synthesis, QA (p=0.008), which is a neurotoxin (86) that is also implicated in ASD etiology (87). NAD^+^ has also been identified as a metabolic link between changes in mtDNA copy number, methionine metabolism and DNA methylation (88). Therefore, both PGC-1α methylation and mtDNA copy number are associated with metabolomic evidence of mitochondrial dysfunction, oxidative stress and neuropathology. Together, our metabolic data are consistent with a dysregulation of the link between methionine metabolism, mitochondrial dysfunction and neurotransmitter synthesis in ASD.

## CONCLUSION

Our study is one of the few molecular studies from Sub-Saharan Africa to examine ASD in an understudied African population. We present the first report of an association between the methylation of genes involved in mitochondrial biogenesis and remodeling, and mitochondrial function in a South African ASD cohort. While our results are correlative and cannot establish causality, they contribute to, and are supported by, the growing body of evidence that point to aberrant DNA methylation and mitochondrial dysfunction in ASD aetiology. Our results highlight the value of epigenetic research in under-studied populations to highlight novel associations. The central role of DNA methylation in modulating mitochondrial function highlights the potential to explore existing mitochondrial medications as putative therapeutic interventions for ASD symptomatology.

## METHODS

### Participants and Sample collection

The ASD vs controls design and study participants were previously described (22). This study examined South African boys with ASD (n=59) and age-matched typically developing controls (n=40) from three demographic groups: African -, European- and Mixed-ancestry. All participants completed an ADOS-2 assessment which was used for phenotyping in the ASD group and ensured the absence of ASD traits in the control group. The study protocol had University of Cape Town Ethics, as well as the Western Cape Government approval to recruit participants at schools. Buccal cells were collected from participants for DNA extraction; this is a minimally invasive collection method suited for DNA methylation studies (89,90). Urine was collected for organic acids extraction for metabolomic analysis using GC-MS.

### Targeted Next-Generation Bisulfite Sequencing

DNA methylation was quantified using tNGBS for PGC-1α and STOML2 in ASD (n=55) and controls (n=44) as well as for FIS1, MFN2, OPA1 and NRF2A in ASD (n=22) and controls (n=22). The tNGBS was completed by EpigenDx, Inc. (MA, USA) who designed a total of 32 tNGBS assays to analyse 171 CpG sites across six genes. They designed seven assays that analysed 26 CpG sites for PGC-1α, eight assays that analysed 45 CpG sites for STOML2, four assays covering 30 CpG sites for FIS1, five assays covering 26 CpG sites for MFN2, five assays covering 25 CpG sites for OPA1 and three assays that analysed 19 CpG sites for NRF2A. Methylation levels were calculated by dividing the number of methylated reads by the number of total reads. Unpaired two-tailed t-tests with unequal variance were used to determine the significant DM CpG sites between ASD and control (p<0.05).

### Mitochondrial DNA copy number and deletion

mtDNA copy number and mitochondrial deletions were measured using multiplex real-time quantitative polymerase chain reaction (RT-qPCR). Mitochondrial genes, ND1 and ND4, were amplified by RT-qPCR and normalized to the nuclear gene, B2M, in the same PCR reaction. The probes were coupled to nonfluorescent quenchers (BHQ®, LGC BioSearch) and both the primers and probes used were previously reported by Grady *et al*. (64). Each DNA sample (20ng/μl) was amplified in triplicate, with standard curves set for each gene using equimolar pooled DNA from ASD and controls in a 10-fold dilution series. A two-step thermal profile was used with denaturation at 95°C for 10 min, followed by 40 cycles of 10s at 95°C, 30s at 60°C on the Rotor-Gene Q 6-plex (QIAGEN). Mitochondrial copy number was calculated using the equation 2 × 2ΔCt, where ΔCt = Ct (nuclear DNA gene) – Ct (mtDNA gene). Significance was determined using a two-tailed unpaired t-test.

### Metabolomic correlation analysis

We examined the correlation between PGC-1α methylation, mtDNA copy number and levels of urinary organic acids associated with mitochondrial dysfunction. Urinary organic acids were previously extracted and quantified by GC–MS for 35 participants (21 ASD and 13 controls) (22). The GC–MS data had been analyzed and deconvoluted using a standard metabolomics-based data processing workflow (75) and was log 2 transformed before statistical analysis. The Shapiro-Wilks test was used to test for normality, after which the Spearman Rank Correlation analysis was performed in SPSS (v26).

## ABBREVIATIONS

ADOS-2: Autism Diagnostic Observation Schedule, Second Edition
ASD: Autism Spectrum Disorder
ATP: Adenosine 5’-triphosphate
BCAAs: Branched-chain amino acids
B2M: Beta-2-microglobulin
CA: Citramalic acid
CREB1: CAMP response binding element 1
DM: Differentially methylated
DRP1: Dynamin-related protein 1
FIS1: Mitochondrial fission protein 1
GABPA: GA-binding protein α-chain
GC-MS: Gas chromatography – mass spectrometry
MFN1: Mitofusin 1
MFN2: Mitofusin 2
mtDNA: Mitochondrial DNA
ND1: NADH dehydrogenase 1
ND4: NADH dehydrogenase 4
Nrf-1: Nuclear respiratory factor 1
Nrf-2: Nuclear respiratory factor 2
NRF2A: Nuclear respiratory factor 2 alpha subunit
OPA1: Optic atrophy 1
OXPHOS: Oxidative phosphorylation
PGC-1α: Peroxisome proliferator-activated receptor gamma coactivator -1 alpha
ROS: Reactive oxygen species
RT-qPCR: Real-time quantitative polymerase chain reaction
STOML2: Stomatin-like protein 2
TFAM: Mitochondrial transcription factor A
TFB2M: Mitochondrial transcription factor B2
tNGBS: targeted Next Generation Bisulfite Sequencing
TSS: Transcriptional start site
UTR: Untranslated region
3-H-3-MGA: 3-hydroxy-3-methylglutaric acid
3-MGA: 3-Methylglutaconic acid

## DECLARATIONS

### Ethics approval and consent to participate

This study was approved by the University of Cape Town (FSREC076-2014) and Western Cape Government (20141002-37506). We obtained informed, written consent from the parents of all the participants prior to their participation in the study.

### Consent for publication

Not applicable.

### Availability of data and other materials

The molecular datasets used and analysed during the current study are available from the corresponding author on reasonable request.

### Authors’ contributions

CO conceptualised the overall study design, was responsible for the phenotype data, supervised the laboratory work and data analysis and was the major contributor in writing the manuscript. These authors contributed equally to the manuscript: SB, EM and CM. SB assisted with the design, data acquisition and analysis of the mitochondrial DNA copy number data and contributed to writing the manuscript. EB assisted with the design and analysis of the methylation data for the mitochondrial fission and fusion genes and contributed to writing the manuscript. CM assisted with the design and analysis of the methylation data for the mitochondrial biogenesis genes, analysed the urinary metabolic data, and contributed to writing the manuscript. All authors read and approved the final manuscript.

## Acknowledgements

We thank the National Research Foundation (Grant No 118524), South Africa for funding this research. We are grateful to the families and the staff of all the schools that participated in our research.

## LIST OF ADDITIONAL FILES

**Additional File 1**.**xls: Supplementary Tables**

**Table S1:** Summary of demographic and phenotypic characteristics of participants used in this study. ASD = Autism Spectrum Disorder; ADOS-2 = Autism Diagnostic Observation Schedule, Second Edition.

**Table S2:** Differentially methylated CpG sites (p<0.05, methylation range > 5%) of PGC-1α and STOML2 (n = 55 ASD, 44 controls) and NRF2A, MFN2, FIS1 and OPA1 (n = 22 ASD, n = 22 controls).

**Table S3A:** Spearman correlation coefficients for urinary organic acids that correlate significantly with mtDNA copy number (n = 20 ASD, n =13 controls). The most significant (p<0.01) correlations with r<-0.3 or r>0.3 are shown in bold, non-significant correlations are not shown. BCAA-Branched Chain Amino Acids, TCA - Tricarboxylic acid cycle.

**Table S3B:** Spearman correlation coefficients for urinary organic acids (n= 20 ASD, n = 13 controls) that correlate significantly with methylation at promoter of PGC-1α at the CpG#1 site.

**Additional File 2**.**png : Fig. S1:** PGC-1α is differentially methylated in ASD. Targeted Next-Generation Bisulfite Sequencing (n = 55 ASD, n= 44 controls) identified differential methylation 12 CpG sites on PGC-1α. Differential methylation is shown as fold change in percentage methylation in ASD relative to Controls.

**Additional File 3**.**png: Fig. S2:** The relationship between PGC-1α promoter methylation and mitochondrial DNA (mtDNA) copy number and deletions is altered in ASD. Heatmap shows significant Spearman rank correlations between mtDNA deletions, mtDNA copy number and percentage methylation at differential methylated CpG sites in the PGC-1α promoter (n = 49 ASD, n = 42 controls);*indicates p<0.05. PGC-1α CpG#1 correlated positively with mtDNA copy number in ASD (Spearman’s r = 0.9, p = 8.814 10-10) while CpG#4 correlated negatively with both mtDNA copy number and (Spearman’s r = −0.4, p = 0.045) and mtDNA deletions (Spearman’s r = −0.4, p = 0.032) in controls.

