## Supplementary figures and images for "DNA methylation of PGC-1α is associated with elevated mtDNA copy number and altered urinary metabolites in Autism Spectrum Disorder"

### Supplemental Fig1

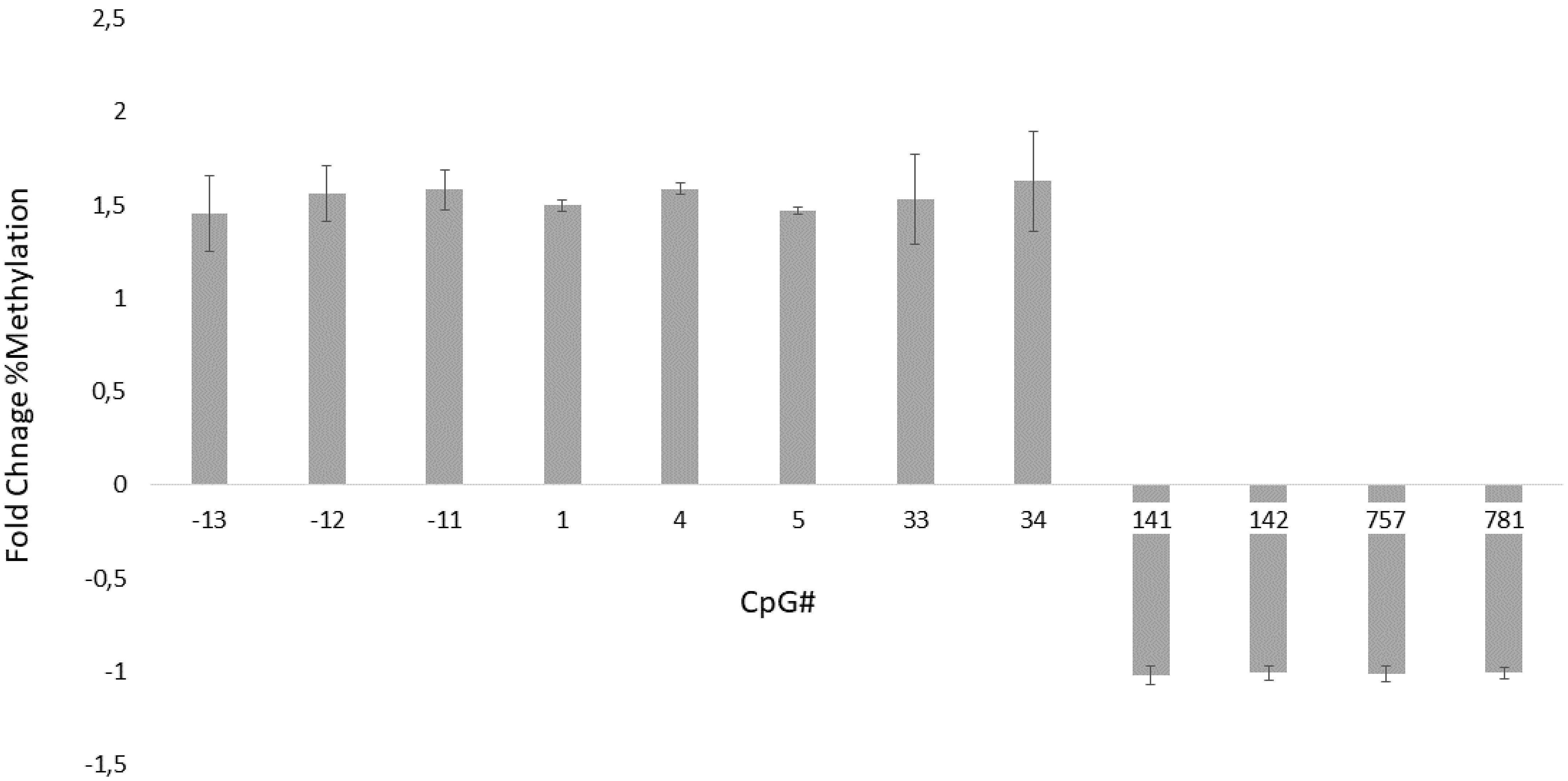

### Supplemental Fig2

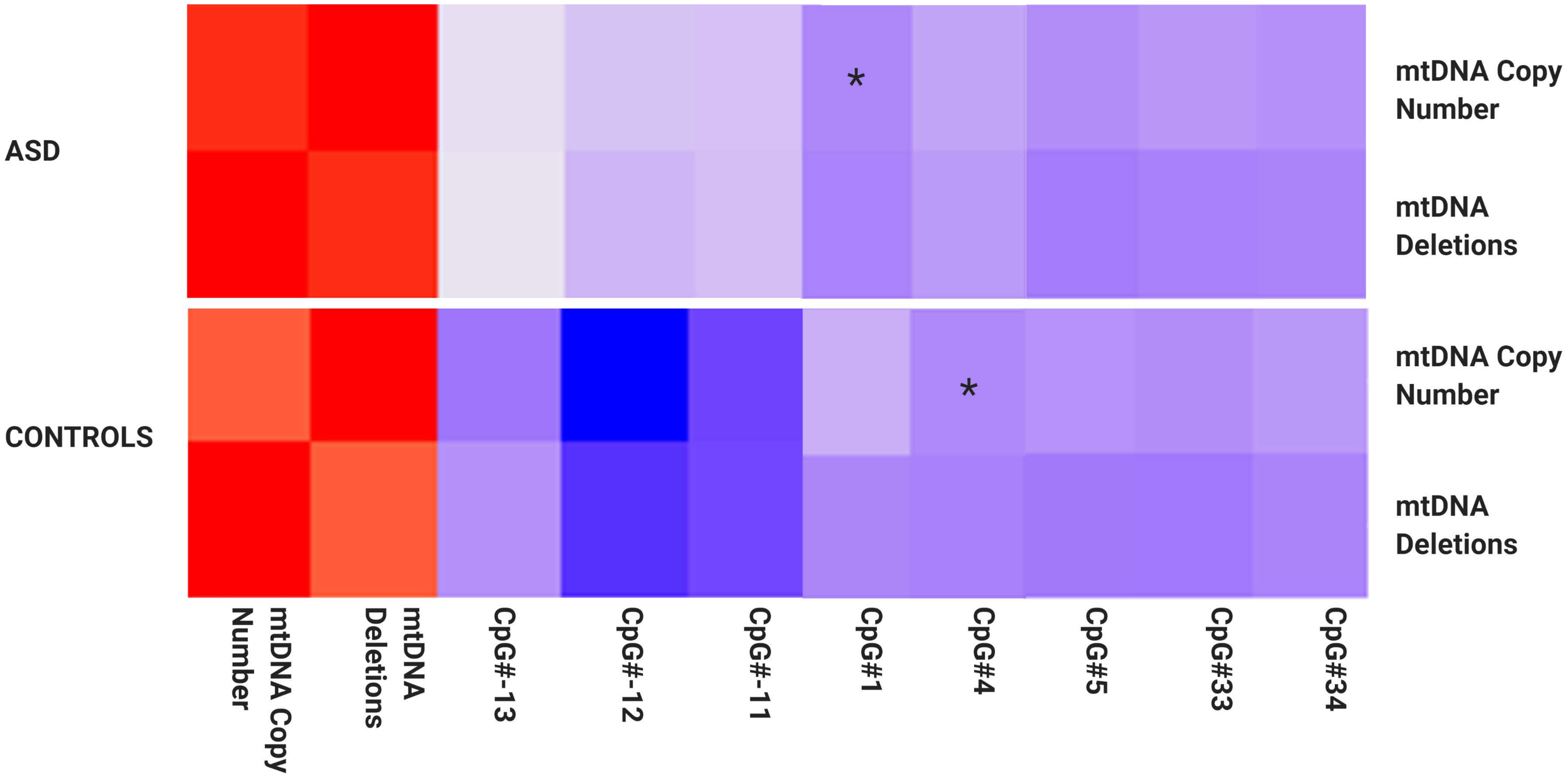
